# Sequence analysis and confirmation of type IV pili-associated proteins PilY1, PilW and PilV in *Acidithiobacillus thiooxidans*

**DOI:** 10.1101/350900

**Authors:** Elvia Alfaro-Saldaña, O. Araceli Hernández-Sánchez, Araceli Soberano-Patrón, Marizel Astello-García, J. Alfredo Méndez-Cabañas, J. Viridiana García-Meza

## Abstract

*Acidithiobacillus thiooxidans* is an acidophilic chemolithoautotrophic bacterium widely used in the mining industry due to its metabolic sulfur-oxidizing capability. The biooxidation of sulfide minerals is enhanced through the attachment of *A. thiooxidans* cells to the mineral surface. The Type IV pili (TfP) of *At. thiooxidans* may play an important role in the bacteria attachment, since among other functions, TfP play a key adhesive role in the attachment to and colonization of different surfaces. In this work, we reported for the first time the confirmed mRNA sequences of three TfP proteins from *At. thiooxidans*, the protein PilY1 and the TfP pilins PilW and PilV. The nucleotide sequences of these TfP proteins show changes of some nucleotide positions with respect to the corresponding annotated sequences. The bioinformatic analyses and 3D-modeling of protein structures sustain their classification as TfP proteins, as structural homologs of the corresponding proteins of *P. aeruginosa*, results that sustain the role of PilY1, PilW and PilV in pili assembly. Also, that PilY1 comprises the conserved *Neisseria*-PilC (superfamily) domain of the tip-associated adhesin, while PilW of the superfamily of putative TfP assembly proteins and PilV belongs to the superfamily of TfP assembly protein. Also, the analyses suggested the presence of specific functional domains involved in adhesion, energy transduction and signaling functions. The phylogenetic analysis indicated that the PilY1 of *Acidithiobacillus* genus forms a cohesive group linked with iron- and/or sulfur-oxidizing microorganisms from acid mine drainage or mine tailings. This work enriches knowledge regarding colonization, adhesion and biooxidation of inorganic sulfurs by *A. thiooxidans*.

## Introduction

*Acidithiobacillus thiooxidans* is an acidophilic chemolithoautotroph that uses reduced sulfurs as a source of electrons and reducing power, including elemental sulfur (S^0^), polysulfides (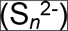) and sulfide minerals, such as pyrite (FeS_2_), chalcopyrite (CuFeS_2_) or sphalerite (ZnS).

Bacterial attachment to mineral surfaces influences the rate of dissolution of the mineral because of surficial phenomena: Mixed potential decreases, changes in kinetics and mass-transport processes [1]. Accordingly, bacterial attachment is due to self-organization by a bioelectrochemical evolution on the interface. Interfacial studies on charge and mass transfer demonstrate that S^0^ biooxidation by *At. thiooxidans* begins in the early stages of interaction (1 to 24 h) when the biofilm is not constituted, and it is primarily controlled by surficial characteristics that pivoted the bacterial attachment to the hydrophobic S^0^; such attachment is an energy-dependent process in which *At. thiooxidans* essentially activates or modifies the reactive properties of S^0^ [2, 3]. The hydrophobic character of the interface “determines the free energy of the adhesion process” [4]. The Type IV pili (TfP) of *At. thiooxidans* may play an important role in the bacterial attachment and bioelectrochemical evolution on the bacteria-mineral interface, *e.g.*, surpassing hydrophilic interactions.

Valdés *et al.* [5] and Li *et al.* [6] suggested that the efficiency of *At. ferrooxidans* to attach and colonize mineral surfaces (*e.g.*, FeS_2_ or CuFeS_2_) and solid reduced sulfur depends on TfP as well as its multiple copies of genes for pili biosynthesis. Other *Acidithiobacillus* species such as *At. caldus* and *At. thiooxidans* contain genes coding for TfP assembly proteins [7–9] that are related to biofilm formation, acting as c-di-GMP effector proteins [10].

The TfP are semiflexible polymeric filaments of pilins anchored to the cellular membrane, of 5-7 nm diameter and from 4-5 μm up to several micrometers in length [11,12]. The TfP have been grouped based on the aminoacidic homology of the pilin subunits, which are relatively conserved in prokaryotes. Pilins share an N-terminal cleavage/methylation (N-methylphenylalanine) domain (NTD) of approximately 25 amino acids (aa) followed by a stretch of hydrophobic residues forming an extended α-helix and a disulfide bond at the C-terminal domain (CTD) [11, 13]. Pilins interact via their conserved NTD α-helix, which forms a hydrophobic core that provides extreme mechanical strength [13]. Further, it has been reported that this conserved NTD in prepilins is also in the type II secretion systems (T2SS), known as pseudopilins [14–17].

Pelicic [18] suggested that the minimal machinery needed for TfP assembly includes pilins and other TfP proteins: (i) Major pilin with NTD (pilin-like motif), (ii) specific peptidase that processes the precursors of pilins or prepilins (*e.g.*, the PilD of *Neisseria* spp.), (iii) traffic ATPase that powers TfP assembly (PilT), (iv) internal inner membrane protein (PilG), (v) integral outer membrane or secretin necessary for the emergence of TfP on the cell surface (PilQ), and (vi) proteins also found in T2SS, the pilin-like proteins.

Interesting, “the TfPs are universal in prokaryotes and have shown extreme **functional** versatility”. Among other **functions** (motility, cell signaling, pathogenic functions, protein secretion, DNA uptake, electrical conductance, and so on), TfP are “sticky organelles” Berry and Pelicic [17] that play a key role as adhesive to stick bacteria to one another and to attach to and to colonize to a wide variety of surfaces, leading to the formation of colonies and even biofilms with cells embedded within an EPS matrix, including “2-D” (thin liquid) and 3-D biofilms [12, 17].

The adhesive ability of TfP is due to the presence of a non-pilin protein, an adhesin (integrin) located on the distal tip of the TfP [17, 19–21]. This adhesin, designated PilC or PilY1, is expressed in multiple species. For instance, PilY1 of *Pseudomonas aeruginosa* is the orthologue of the meningococcal PilC of *Neisseria gonorrhoeae*, and both proteins Although PilY1 and PilC share partial sequence homology, they have a high degree of structural similarity [22, 23]. PilC is a phase-variable protein that belongs to a Conserved Protein Domain Family of several PilC protein sequences [23]. Further, PilC/PilY1 is dispensable for TfP assembly at early stages of TfP biogenesis, for instance, in the absence of pilus retraction [17, 22–24]. Our previous analysis on,showed that the tip-associated adhesin PilY1 of *At. thiooxidans* Licanantay (WP_031573362] exhibited 55% and 86% identity with the type IV pilin biogenesis protein of *At. thiooxidans* ATCC 19377 (WP_010638975.1) and of *At. albertensis* (WP_075322776.1), respectively.

Other proteins involved in adhesion are the minor but highly conserved pilin proteins PilW and PilV of *Neisseria* spp. [17, 24, 25]. The outer-membrane lipoprotein PilW participates in pilus biogenesis for the stabilization of the pilus but not for their assembly, as well as to allow bacterial adherence [26]. Another possible function for PilW is the transfer of electrons, as has been proposed for *At. ferrooxidans* [27] and other species with TfP [28].

Thus, the putative proteins PilY1, PilW and PilV of *At. thiooxidans* were examined in this work. We chose these proteins because of PilY1’s possible function as an adhesin and the role of PilW and PilV as pilus assembly proteins (structural pilins) of the TfP of *At. thiooxidans.* Most genes of *At. thiooxidans* are annotated in GenBank. In this study, the sequences of the TfP proteins PilY1, PilW and PilV were first confirmed and uploaded to GenBank. In addition, bioinformatic analyses and 3D modeling of each TfP protein were performed.

## Materials and methods

### *Acidithiobacillus thiooxidans* culture and maintenance

*At. thiooxidans* strain ATCC 19377 was cultured in ATCC 125 medium (in g/L: 0.4 (NH_4_)_2_SO_4_, 0.5 MgSO_4_7H_2_O, 0.25 CaCl_2_, 3 KH_2_PO_4_, 0.005 FeSO_4_7H_2_O, and 10 S^0^, pH 2.0 adjusted with concentrated H_2_SO_4_). Cultures were aerobically incubated at 29 ± 1 °C under orbital agitation (110 to 120 rpm) for 5, 10, 15 and 21 days.

### Multiplex PCR and cDNA sequencing of *pilYI*, *pilW*, and *pilV*

We used the *At. thiooxidans* ATCC 19377 T draft genome AFOH01000001 originally described by Valdés et al. [7]. The annotated protein sequence of PilY1, PilW and PilV as well as the 16S ribosomal subunit with the accession numbers WP_010638975.1, WP_010638981.1, WP_010638979.1 and AJ459803, respectively, were. Primers were designed against conserved regions of the putative sequences of *pilYI* (ATHI0_RS0106065), *pilW* (ATHI0_RS0106075) and *pilV* (ATHI0_RS0106080) using the software Primer3 [29] and Vector NTI (InforMax-Invitrogen, USA). As a positive control for each PCR reaction, we designed a pair of primers that amplified the 16S rRNA.

Samples for RNA extraction were extracted after 5, 10, 15 and 21 days. At these timepoints, cells were collected by centrifugation at 0.08 × g to separate the cells from S^0^; cells were concentrated and washed by centrifugation at 21.1 × g for 1 min using saline phosphate buffer (PBS; in g/L: 8 NaCl, 1.44 Na_2_HP0_4_, 0.24 KH_2_P0_4_ (J.T. Baker, USA), and 0.2 KCl (Sigma-Aldrich, USA); pH 7.4).

RNA extraction was performed using TRIzol (Invitrogen, USA) reagent according the manufacturer’s protocol. After quantification using a NanoDrop 2000c (Thermo Scientific, USA), cDNA synthesis was performed according to the manufacturer’s recommendation, using 200U M-MLV (Invitrogen, USA) in a 20 μL total volume reaction. The amplification products were purified using the QIAquick kit (Qiagen, USA) following the recommended procedure, quantified and sequenced. The products were sequenced by the Sanger method at LANBAMA (National Laboratory of Biotechnology, San Luis Potosí, Mexico) and LANGEBIO (National Laboratory of Genomic for the Biodiversity, Guanajuato, México) facilities. Each sequence was confirmed at least five times by analyzing amplification products obtained from different culture replicates. The generated sequences were compared with the annotated sequences of each corresponding gene using the Multalin software [30].

### In silico (bioinformatic) analyses of proteins

The confirmed sequences of PilY1, PilW and PilV were analyzed to search for domains, families and functional sites with bioinformatic tools, including Simple Modular Architecture Search, SMART [31], PROSITE [32], and EMBL-EBI [33] and the CDD/SPARCLE domain analysis function [34]. The protein structure homology-modeling was achieved by using the server SWISS-MODEL [35]. The 3-D model of each protein was generated in I-TASSER [36, 37]]. The analyses of possible ligands were performed in the Ligand-Protein Binding Database, BioLip [38].

### Phylogenetic relationships

Phylogenetic relationships were analyzed for the confirmed nucleotide and protein sequences of PilY1, PilW and PilV from *At. thiooxidans* ATCC19377 against those of the NCBI database using BLASTN 2.8.0+ [39, 40], and UniProt [41] bioinformatic resources.

The sequences obtained from NCBI were aligned with those obtained in this work using MEGA7 software [42] by the Neighbor-Joining method. Thus, the evolutionary history was inferred by using the Maximum Likelihood method (ML) based on the Jones-Taylor Thornton (JTT) model [43]; for the bootstrap consensus tree, 500 replicates were performed [44]. The percentage of trees in which the associated taxa clustered together is shown next to the branches. Initial trees for the heuristic search were obtained automatically by applying Neighbor-Join and BioNJ algorithms to a matrix of pairwise distances estimated using a JTT model; the tree with the highest log likelihood is shown. All positions with less than 95% site coverage were eliminated.

### Transmission electron microscopy (TEM) analyses

After 4 days of incubation, *A. thiooxidans* cells were negative stained with 1% phosphotungstic acid (PTA) or 2% uranyl acetate. One aliquot was directly used, and the other was fixed previously to reduce the insoluble S^0^ present in the sample and stabilize pili. Both samples were washed three times at 1500 rpm for 5 min and stained with PTA. The pili were observed in TEM (JEOL 200 CX, Japan) at 100 kV.

## Results and Discussion

### TEM analyses

After the microscopic analyses, we identified pili in all tested conditions (Fig 1a): i) Negative staining with 2% uranyl made the flagellum, pili, and S^0^ visible; ii) Upon negative staining with 1% PTA, the pili and flagellum were surrendered by extracellular polymeric substance; iii) *A. thiooxidans* were previously washed to remove S^0^ from the medium, then stained with 1% PTA, although S^0^ was eliminated, and many flagella and pili were lost as well. We fixed the bacteria before washing, and in this condition, we preserved the pili, arranged in a pili network (iv, insert).

**Figure 1.**
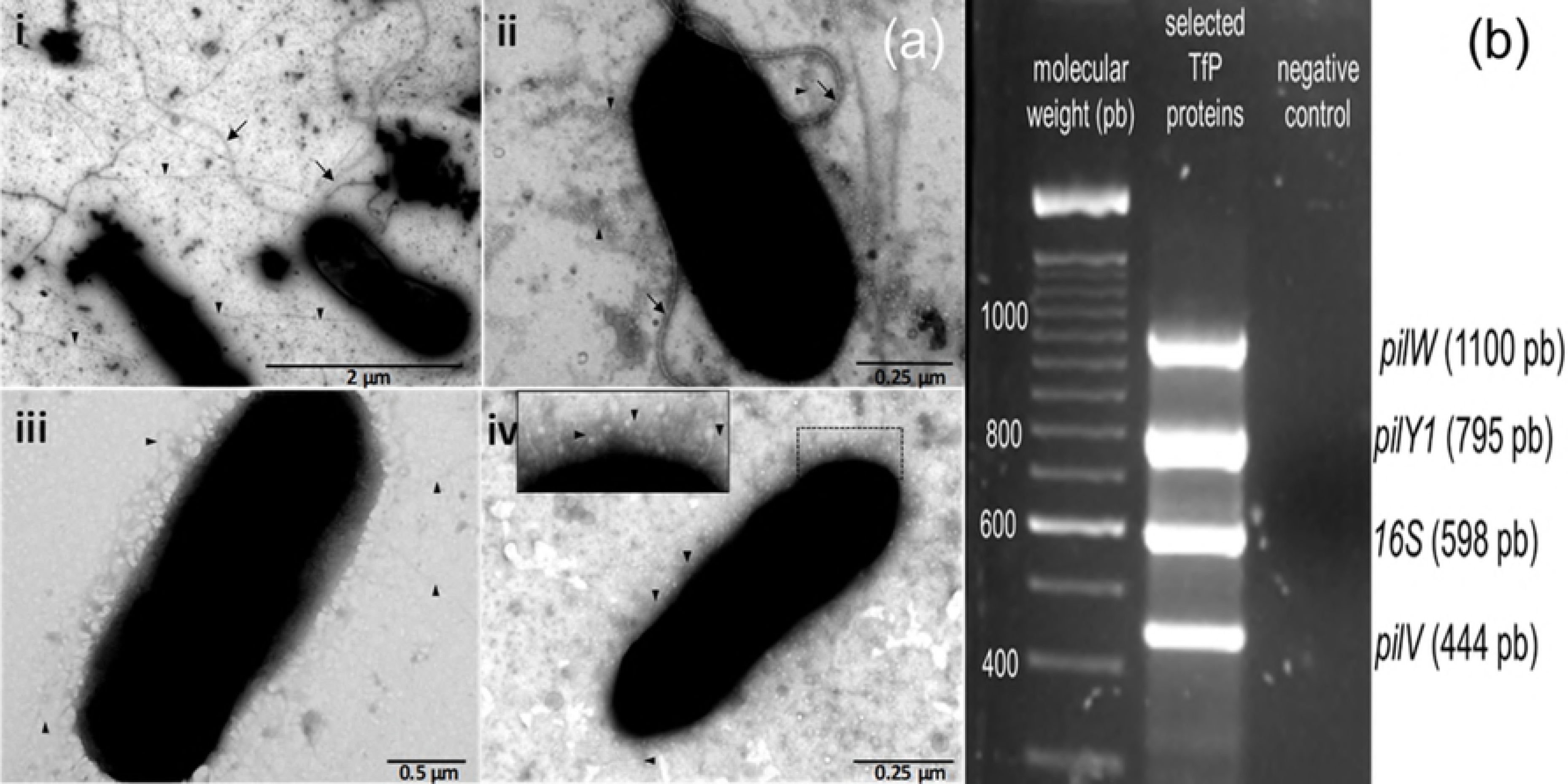
Representative micrographs and multiplex PCR amplification products of *pilYI*, *pilW*, and *pilV*. The TEM microphotographs of *At. thiooxidans* showed the pili (triangles) and flagella (arrows) (**a**). The multiplex PCR was obtained after 5 days of culture; the positive control corresponds to a region of 598 bp of the rRNA 16S (**b**).

### Sequence analyses

For the sequencing of *pilY1*, *pilW* and *pilV*, primers were designed against conserved regions of the putative sequences of 16S from *At. thiooxidans* (GenBank NZ_AFOH01000047.1; [7]. Due to its size (3499 bp), we identified conserved regions of the annotated sequence of *pilY1* comparing it with its homologue in *At. ferrooxidans* strain ATCC 53993 (WP_064218310.1), according to the BLAST analysis. The eight pairs of primers designed (*pilY-1* to *pilY-8*; S2 Table) were used to amplify, purify and sequence (per triplicate) each PCR product of such regions; each obtained amplicon was aligned against the annotated sequence of the *pilY1* gene (ATHIO_RS0106065) to highlight changes between them. After analyzing each sequenced amplicon, the complete sequence was rejoined, and new pairs of primers were designed (*pilY-9* to *pilY-11*, S2 Table) to assess if the highlighted changes were *bonafide*; each pair of these three primers initiates just in the regions with the most significant insertions and deletions previously found. The new obtained amplicons were individually aligned against their corresponding fragment of the reassembled sequence and then with the annotated sequence of *At. thiooxidans*.

A similar strategy was performed for the sequencing of *pilW* and *pilV*, using 5 and 4 pairs of designed primers, respectively (S2 Table); the obtained amplicons were compared with the corresponding annotated sequence of *pilW* (ATHI0_RS0106075) and *pilV* (ATHI0_RS0106080)

Once the complete sequences were obtained, the expression of *pilY1* (MH021598.1), *pilW* (MH021599.1) and *pilV* (MH021600.1) was evaluated at different culture times (Fig 1b).

Thus, the nucleotide sequences of *pilY1*, *pilW* and *pilV* show changes at some nucleotide positions with respect to the corresponding annotated sequences (S1 Table). The changes are more significant for *pilY1* mainly within the N-terminal 1560 bp region. In this region, the difference between the annotated and the confirmed sequences is approximately 25% due to changes of some bases as well as insertions and deletions of 25 and 7 nucleotides, respectively. In the second half of the sequence (1991 bp), the differences between both annotated and confirmed sequences is just 3% due to two insertions of six nucleotides in the confirmed sequence.

However, annotated and confirmed protein sequences showed a pairwise distance of 0.13 (Fig 2a). The PilY1 (AWP39905.1) sustains 85% identity (1007 identical position) with the annotated WP_010638975.1 reported in GenBank. According to BLAST analysis, PilY1 (AWP39905.1) sustains 99% and 77% identity with the sequence of the type IV pilin biogenesis protein of *At. thiooxidans* (WP_065968128.1) and the hypothetical protein of *At. ferrooxidans* (WP_064218310.1); these sequences comprise the *Neisseria-PilC* superfamily domain [34].

**Figure 2.**
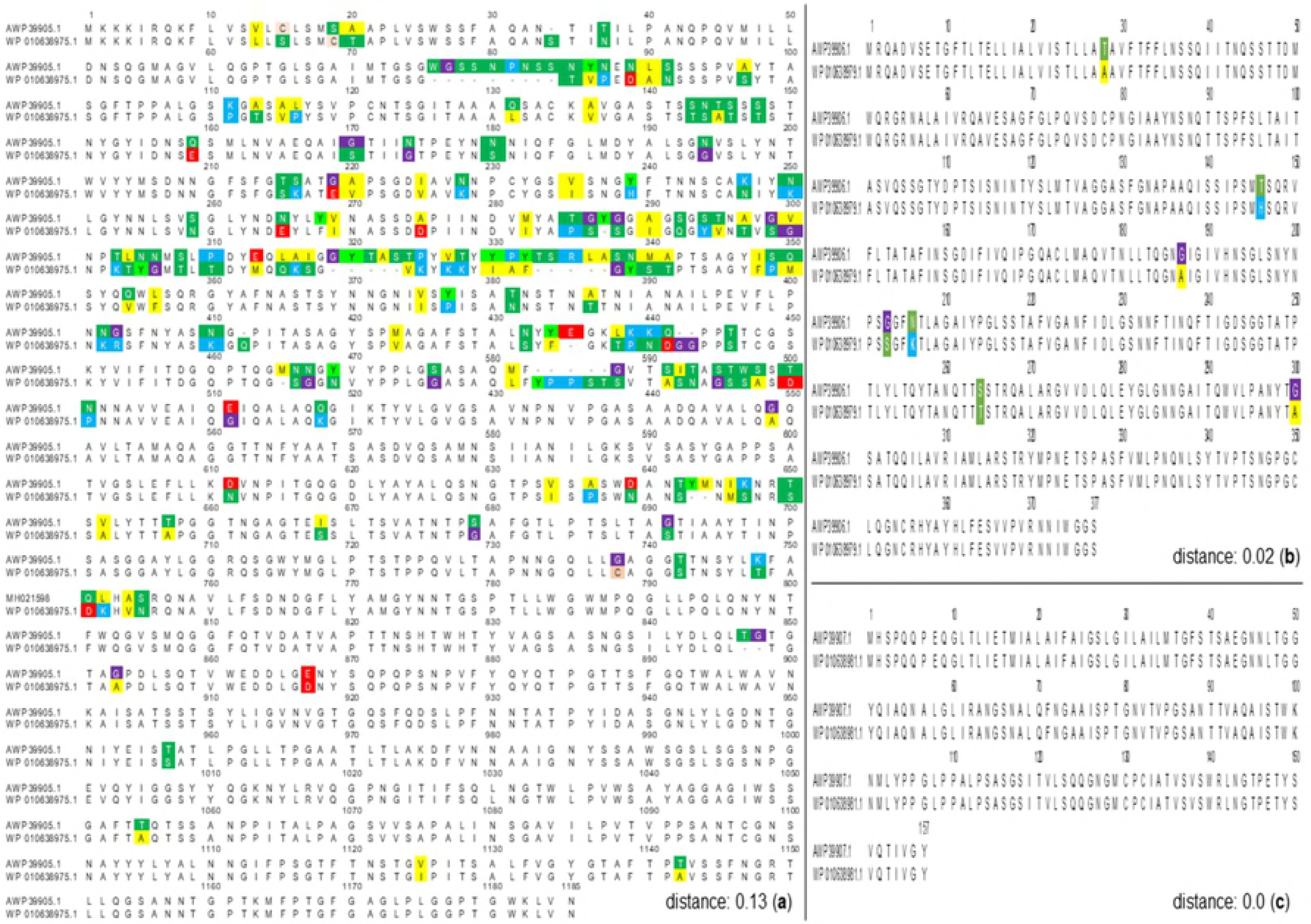
Comparation between the confirmed and annotated sequences. (**a**) PilY1 AWP39905.1 and WP_010638975.1, (**b**) PilW AWP39906.1 and WP_010638979.1 and (**c**) PilV AWP39907.1 and WP_010638981.1, by multiple sequence alignment accuracy and high throughput (MUSCLE) and toggling conserved sites at 100% level (in colors); computed pairwise distance (Poisson model) is also shown. Analyses were done in MEGA7 [42].

Although *pilW* (MH021599.1) shows 20 synonymous changes, the protein sequence (AWP39906.1) shows 98% identity against the annotated and translated sequence of the prepilin-type N-terminal cleavage/methylation domain-containing protein, WP_010638979.1 (ATHIO_RS0106075; Fig 2b). Both sequences comprise a region of 132 aa, the PilW superfamily of putative TfP assembly proteins, as well as *At. ferrooxidans* ATCC 23270 (WP_064218312.1; 83% identity with *At. thiooxidans* AWP39906.1).

Finally, the *pilV* sequence of (MH021600.1) exhibited three synonymous changes that resulted in an identical protein to the annotated and translated WP_010638981.1 (Fig 2c). PilV (AWP39907.1) belongs to the PilV super family of TfP assembly protein, an extracellular structure involved in cell motility [34]. Blast analysis for PilV (AWP39907.1) indicated 92% identity with the pilin (putative) of *At. ferrooxidans* ATCC 23270 (ACK80286.1).

### Proteins bioinformatic analyses and phylogenetic trees PilY1

The bioinformatic analyses of PilY1 (AWP39905.1) confirm that it is a non-pilin protein, essentially hydrophilic (gravy −0.081), and its last 462 aa (48.19 kDa region 715-1176 aa) shares sequence and structural homologies with the C-terminal domain (CTD) of PilY1 of *Ps. aeruginosa* (3hx6.1) [23] that belongs to the conserved domain *Neisseria*-PilC superfamily [34]. In contrast, the NTD of PilC of *Neisseria* spp., PilY1 of P. *aerugionosa* and the PilY1 reported in this work are non-conserved, divergent sequences. Both NTD and CTD sequences of the *At. thiooxidans* PilY1 share similarities with the TfP biogenesis protein of *Acidithiobacillus* spp. (Fig 3).

**Figure 3.**
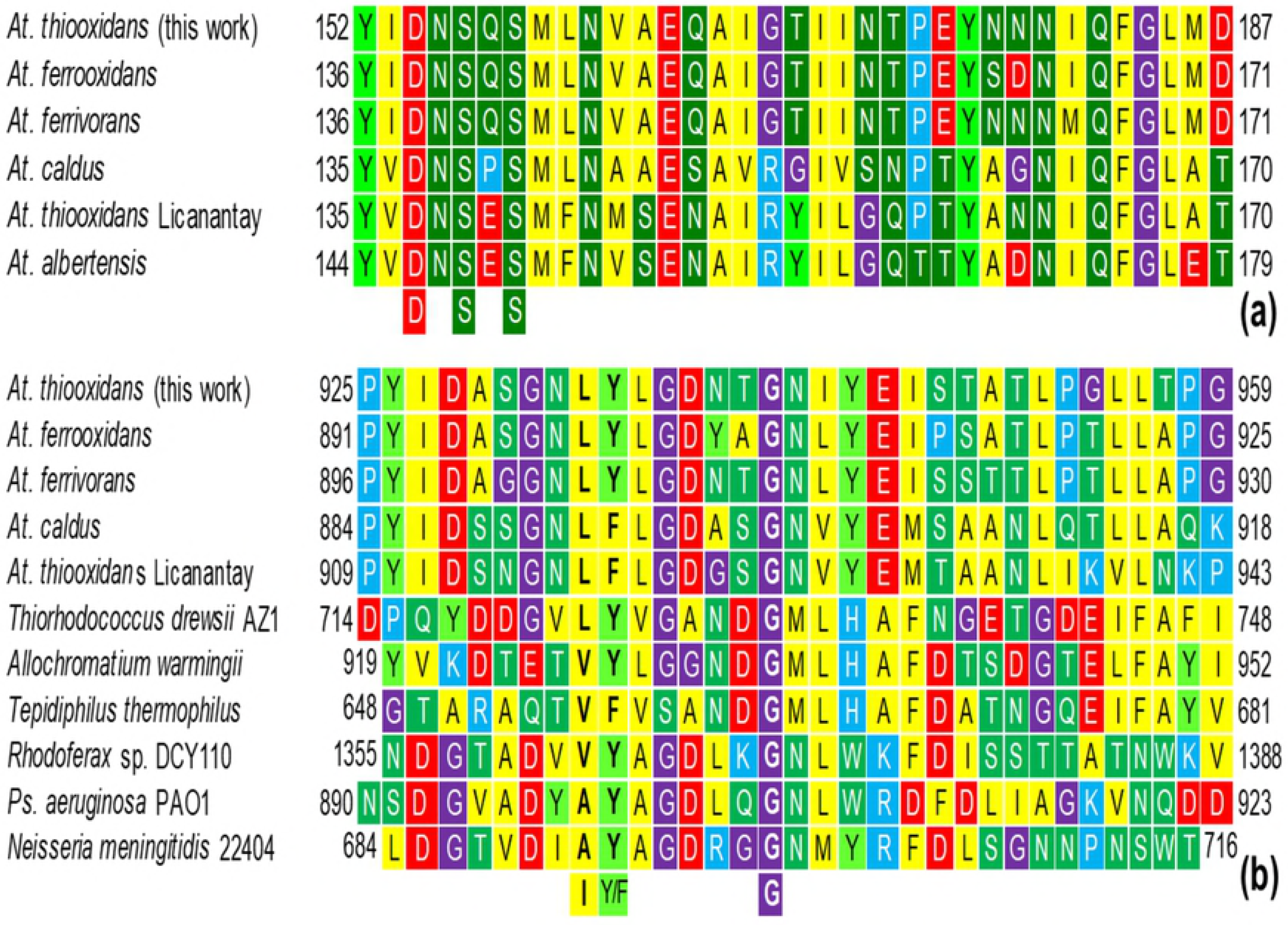
Multiple sequence alignment of PilY1 (AWP39905.1) of *At. thiooxidans*. Showing two regions: (**a**) part of the vWA domain within the NTD of some *Acidithiobacillus* spp. showing the MIDAS (DxSxS) motif, and (**b**) the PQQ domain with the functional motif LYxxxxxG within the CTD of some *Acidithiobacillus* spp. and other bacteria with PilY1/PilC.

The previous is evidenced in the phylogenetic tree (Fig 4), wherein the *Acidithiobacillus* genus forms a cohesive group, deeply related with iron- and/or sulfur-oxidizing microorganisms from acid mine drainage (AMD) or mine tailings such as *Th. bhubaneswarensis*, *Ac. thiooxydans*, *Ga. acididurans*, *Sulfuriferula* sp., and those of the AMD metagenome [45–49]. This phylogenetic tree also reveals that PilY1 and PilC are homologues; all the analyzed sequences comprise the *Neisseria*-PilC beta-propeller domain. Thus, the phylogenetic analysis of PilY1/PilC suggests homologating the nomenclature of this TfP protein (Fig 4), perhaps as the *Neisseria* spp. pilus assembly/adherence protein PilC.

**Figure 4.**
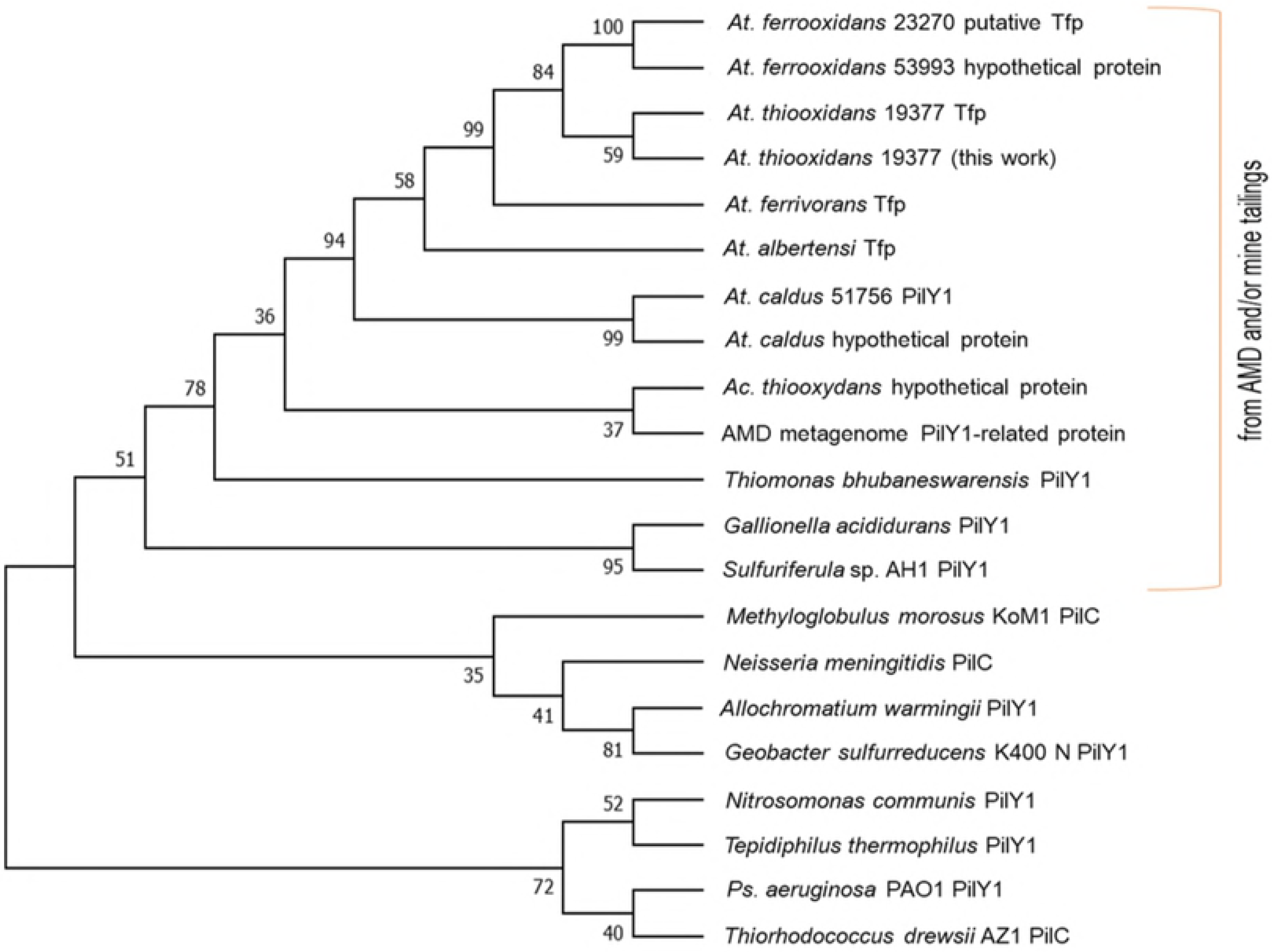
Molecular phylogenetic analysis of PilY1 (AWP39905.1). The evolutionary history obtained by ML (log likelihood: −8108.55) of PilY1/PilC, using 21 aa sequences and the PilY1 (AWP39905.1). There were 212 positions in the final dataset. Evolutionary analyses were conducted in MEGA7 [42]. The aa sequences used for alignment and phylogenetic tree were derived from: acid mine drainage (AMD) metagenome (CBI05240.1), *A. warmingii* DSM 173 (SDX64909.1), *Acidifferobacter thiooxydans* ZJ (OCX45123.1), *At. albertensis* DSM 14366 (WP_075322776), *At. caldus* ATCC 51756 (AIA55059.1 and WP_064306242.1), *At. ferrivorans* YL15 (WP_085537817.1), *At. ferrooxidans* ATCC 23270 (ACK78377.1) *At. ferrooxidans* ATCC 53993 (ACH83324.1), *At. thiooxidans* (WP_010638975.1) *At. thiooxidans* Licanantay (WP_031573362.1), *Ga. acididurans* (isolate ShG14-8; KXS31914.1), *Ge. sulfurreducens* KN400 (ADI84872.2) *Methyloglobulus morosus* KoM1 (ESS74050.1); *Ns. meningitidis* (WP_101069668.1), *Nitrosomoas communis* Nm110 (SDW55994.1), *Ps. aeruginosa* PAO1 (AAA93502.1), *Rhodoferax* sp. DCY110 (WP_076201095.1), PilC of *Sulfuriferula* sp. AH1 (WP_087447088.1), *Te. thermophilus* JCM 19170 (CUB07558.1), *Th. bhubaneswarensis* DSM 18181 (CUA94317.1), *Thiorhodococcus drewsii* AZ1 (EGV31806.1). Tfp: type IV pilin.

Our results are consistent with the predicted 3-D model of *At. thiooxidans* PilY1 (AWP39905.1; Fig 5). This model was generated by I-TASSER based on structures from the PDB (5j44A, 3hx6, 6emkA, 3ja4A and 3iylW), highlighting the PilY1 CTD structure of *P. aeruginosa* (3hx6A and 3hx6) [23]. According to the homology-modeling, PilY1 of *At. thiooxidans*(AWP39905.1) showed similarity (22.73%) with such PilY1of *P. aeruginosa*, mainly among the CTD that corresponds to the *Neisseria-PilC* beta-propeller domain of the tip-associated adhesin PilY1.

**Figure 5.**
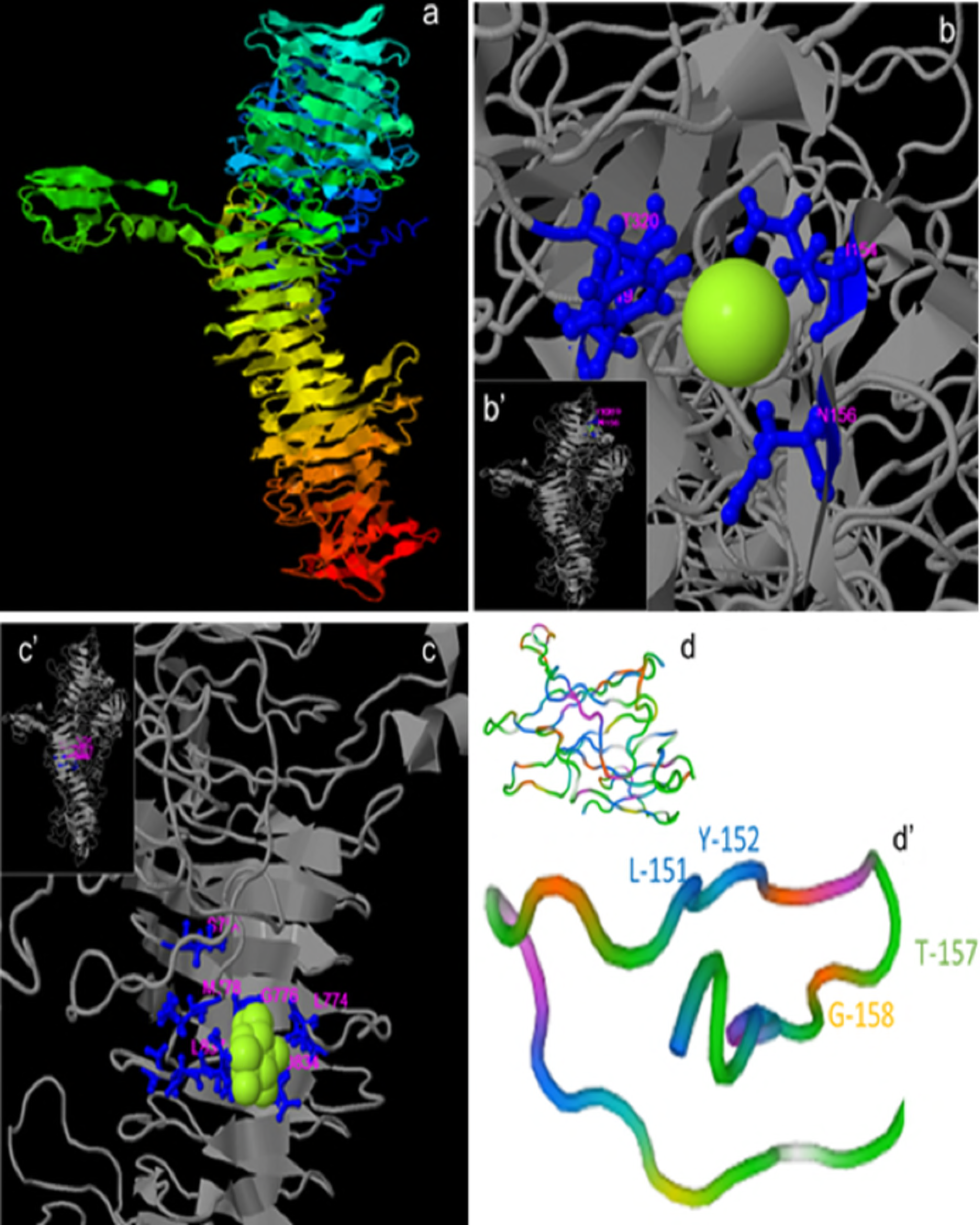
3-D models of the non-pilin protein PilY1 (AWP39905.1). The model shown has a TM-score of 0.61±0.14 with *Ps. aeruginosa* (TM-score >0.5 indicates a correct topology model, while TM-score <0.17 indicates a random similarity) **(a)**. A region of vWA showing the binding ligands L156, Y156, Y-319 and T-320 of MIDAS, for Ca^2+^ binding (**b** and **b**’). a region CTD of PilY1/PilC (TM-score of 0.62±0.14) showing the binding residues for carbohydrates (*e.g.*, α-D-glucose, β-glucose, α-dextrose) by S-754, L-774, G-776, M-778, D-834, L-835, Q-836, L-837 (**c** and **c**’); 3-D modeling conducted in I-TASSER [36, 37] and BioLip [38]. Model of the PQQ region predicted by homology [35] and the anchor motif LYxxxxTG (**d** and **d**’).

*At. thiooxidans* PilY1 (AWP39905.1) bioinformatic analysis confirmed the presence of five regions or motifs (Figs 3 and 5) also found in TfP proteins from *At. ferrivorans* SS3 (WP_085537817.1), *At. ferrooxidans* ATCC 53993 (ACH83324.1), *At. albertis* (WP_075322776), *At. caldus* MTH-04 (WP_064306242.1), and *At. thiooxidans* Licanantay (WP_031573362):

I. A von Willebrand factor type A (vWA) domain (540 aa) was found in the region from amino acid 42 to 581, folded into a α/β Rossmann fold (alternating β-strand with α-helix). Cellular functions such as cell migration, adhesion and signaling have been associated with the vWA domain [34, 50]. E-value (BLAST): 1×10^−28^.
II. Within the vWA, a metal ion-dependent adhesion sites (MIDAS motif, for Mn^2+^, Mg^2+^ or Ca^2+^) of 35 aa residues (region 153-187) folded as a short coil followed by an α-helix. MIDAS are commonly present in cell surface-adhesion receptors or molecules (CAMs), *e.g.*, integrins [51]; thus, such components are involved in cell-cell and cell-matrix interactions through its adhesion function. MIDAS comprised the conserved components DxSxS, T, and D (Fig 3a) [34, 52]. Modeling by homology [35], the MIDAS motif of *A. thiooxidans* PilY1 (AWP39905.1) exhibited 34.38% similarity (covering 91%, from 2 to 33 aa) with the model of a cell wall surface anchor family protein (PDB: 3tw0.1.A), specifically with the adhesive tip pilin of *Streptococcus agalactiae* [50]. The constructed 3-D model specifies that the MIDAS motif within vWA includes two different loops (α-helix) exposed in the protein surface (Fig 5b and b’), a location that allows it to interact with divalent metal ions such as Ca^2+^ via the binding ligands I-154, N-156 and T-320 of MIDAS.
III. The aforementioned conserved CTD or *Neisseria*-PilC domain is 462 aa (715-1176 aa) mainly composed by aliphatic aa (G, A, V, L and S, T; 60.2%) and slightly hydrophilic (gravy:-0.094). The 3-D model of this CTD (Fig 5c) was based only on structures of the CTD of *P. aeruginosa* PilY1, 3hx6 [23], showing a normalized Z-score of the threading alignments, up to 7.52. Other structural analogs predicted using I-TASSER, are oxidoreductases-substrate oxidation and electron transfer (PDB: 1h4jE, 1lrwA, 2d0vI, 1yiqA1, 1FLG), with Ca^2+^ or pyrroloquinoline quinone (PQQ) as ligands (see below).
IV. Within the *Neisseria*-PilC domain, the 3-D model also predicts binding sites for carbohydrates such as glucose (Fig 5c), previously described for tail-spike proteins for recognition and adhesion of *Salmonella* and *E. coli* bacteriophage HK620 [53].
V. Also within the *Neisseria*-PilC domain is a motif that occurs in propellers of PQQ cofactor binding domains (SMART and InterPro accession numbers SM00564 and IPR018391, respectively) of 27 aa (region 926-952) that catalyzes redox reactions; *e.g.*, a β-propeller repeat occurring in enzymes with PQQ as cofactor in prokaryotic quinoprotein dehydrogenases that are involved in electron transfer processes, and thus energy transduction [54]. *Ps. aeruginosa* PilY1 (AAA93502) and PilC of *N. meningitidis* 22404 (WP_101069668) also include the PQQ cofactor binding domain, while most of the *Acidithiobacillus* spp. and other sulfur-oxidizing microorganisms comprise the anchor motif LYxxxxTG (Figs 5b and d). The iron- and/or sulfur-oxidizing microorganisms from AMD (e.g., *Th. bhubaneswarensis*, *Ac. thiooxydans*, *Ga. acididurans*, *Sulfuriferula* sp.; Fig 4) also contain the PQQ domain (Fig 3b), while the PilY1-related protein (fragment) from the AMD metagenome (CBI05240.1) mostly corresponds to the vWA domain within the NTD that also comprises the MIDAS motif, DxSxSxxxxxxxxxxT (Fig 3a). Further, the other sulfur-oxidizing microorganisms presented in the phylogenetic tree (Fig 4), *Te. thermophilus* and *T. drewsii* AZ1, included the PQQ domain in their tip-associated adhesin PilY1/PilC, according to SMART analyses [31]. The presence of PQQ in such PilY1/PilC proteins suggests that this adhesin initiates the biooxidation of reduced compounds (*e.g.*, S^0^ or metal sulfides as FeS_2_, and CuFeS_2_) and may be involved in electron transfer between the substrate and other components. *Sensu* Li and Li [27], the biooxidation of Fe(III) by *At. ferrooxidans* depends on functional pili which transfer electrons from the reduced Fe to the cell surface, while *At. ferrooxidans* attaches strongly to solid surfaces such as FeS_2_ [6].

### Pilins PilW and PilV

The 3D-modeling of PilW and PilV of *At. thiooxidans* (Figs 6a and b) revealed high similarities to the pilin of *Ps. aeruginosa* strain K, PAK (PDB key: 1oqw). For PilW, the model was also based on structures of a protein with structural similarity to flagellin of *Burkholderia pseudomallei* (4ut1A), a putative peptide-binding domain (adhesin) of *Marinomonas primoryensis* (5k8g), and the pilin FimA for adhesion from *Dichelobacter nodosus* (3sok).

**Figure 6.**
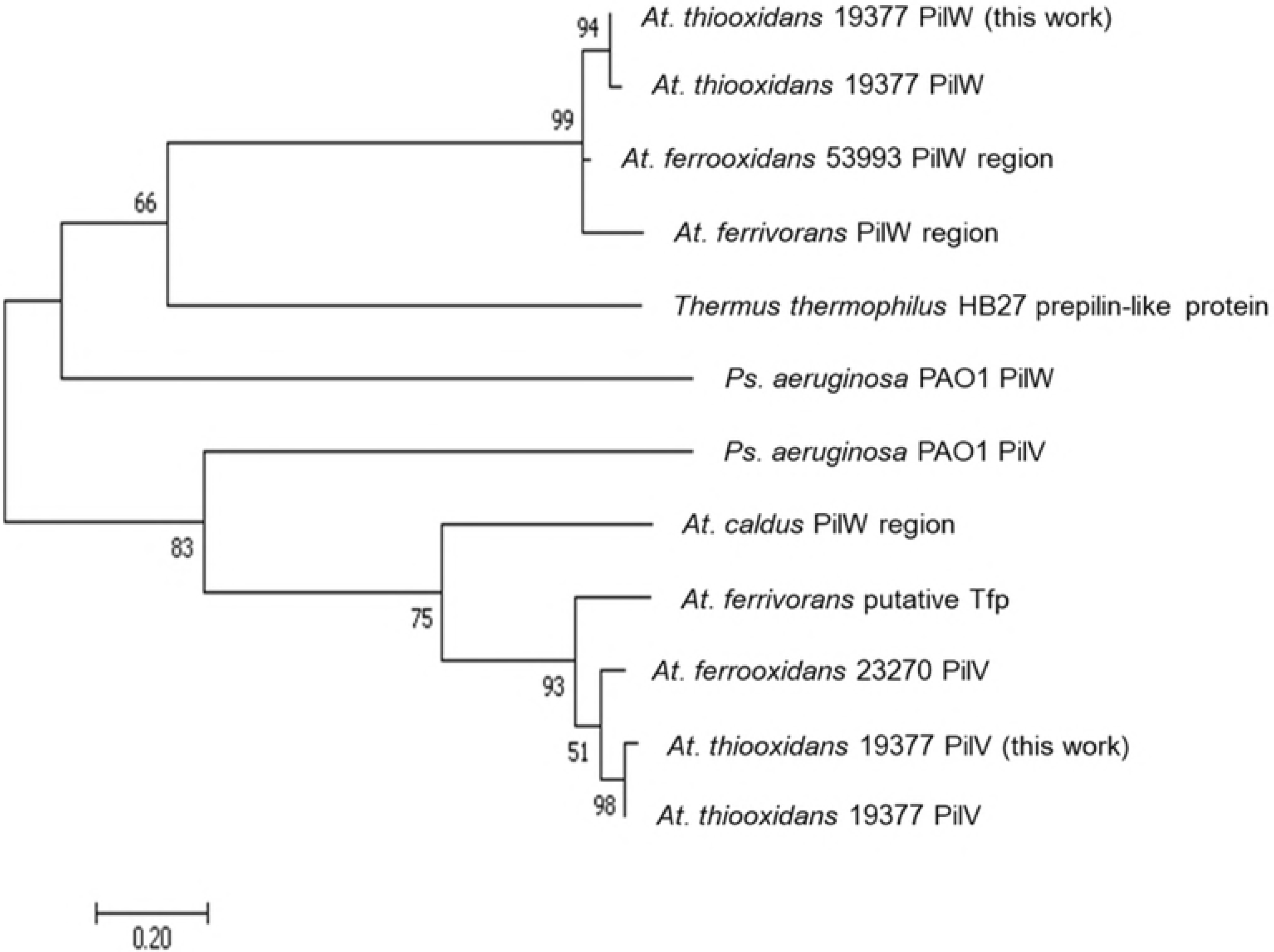
Bioinformatic analyses of the confirmed sequences of *At. thiooxidans* PilW (AWP39906.1) and PilV (AWP39907.1). (**a**) 3-D model of PilW (TM-score: 0.42±0.14) and (**b**) PilV (TM-score: 0.49±0.15) conducted in I-TASSER [36, 37] and BioLip [38]; the models were obtained based on structures from the PDB data base; for PilW: 5k8g, 1oqw, 4ut1A, 3sok, 4m00, 5gaoE; for PilV: 1oqw, 3sok, 5bw0, 1ay2 and 3ci0. Modeling by homology of the NTD of (**c**) PilW (1-81 aa) and (**d**) PilV (11-62 aa) using as template the 3nje.1 minor pseudopilin from the *Ps. aeruginosa* [15]. for PilW, and the 5vxy.1.E pilin of *Ps. aeruginosa* and *N. gonorrhoeae* [55, 56] for PilV. (**e**) Multiple sequence alignment conducted in MEGA7 [42], of the region NTD (25-28 aa) of the confirmed PilW (AWP39906.1) and PilV (AWP39907.1) sequences of *At. thiooxidans*, and of other sequences from the gene bank of prepilin containing proteins of *At. thiooxidans* ATCC 19377 (WP_010638979.1), *At. thiooxidans* Licanantay (WP_051690664.1). *At. ferrooxidans* ATCC 593993 (ACH83326.1), *At. ferrivorans* (WP_035191506.1), *At. albertensis* (WP_075323115.1) and *At. caldus* (WP_070113768.1), as well as TfP pilins or prepilins of *Ps. auriginosa* PAO1 (NP_253215.1) *N. gonorrehaea* (SBO76855.1), *E. coli* (PIM50236.1), *Microcystis aeruginosa* (WP_012267210.1), *Desulfovibrio magneticus* (EKO40131.1), *Shewanella baltica* (WP_012588380.1) and *Mariprofundus micogutta* (WP_083530569.1).

As the sequences 1oqw and 3sok, PilW and PilV of *At. thiooxidans* and other TfP pilins have leader peptides of approximately 20-25 aa in the NTD with a highly conserved G and the GFXXXXE domain (Fig 6e). According to Dalrymple and Mattick [57] and Mattick [58], all the necessary information for the processing of subunits and assembly of pili is in these first aa of the α1-N; thus, mutations of NTD residues can markedly affect pilus assembly [13]. Specifically, the conserved glutamic acid E5 is the only charged residue in NTD (Fig 6b’), which is essential for pilus assembly and seems to be required for the methylation step [34, 55, 59]. In *At. thiooxidans*, E5 interacts with divalent cations such as Ca^2+^ (Fig 6b).

The CTD of PilW includes a region from aa residue 239 to 371 that is identified as the “TfP pilus assembly protein PilW” (pfam16074). Its predicted secondary structure is mainly strands and coil, that define a globular head (Fig 6a), a head also observed in the 3-D model of PilV (Fig 6b). This globular head domain is a structurally variable region among the different pili, variability that imposes different functions, *e.g.*, specific subunit interactions and packing arrangements within the filament, pilus-pilus interaction, and interactions between the pili and their environment [55].

The molecular phylogenetic analysis confirms that PilW and PilV of *At. thiooxidans* are core subunits of TfP, similar to PilW and PilV of *Ps. aeruginosa* PAO1 (Fig 7).

**Figure 7.**
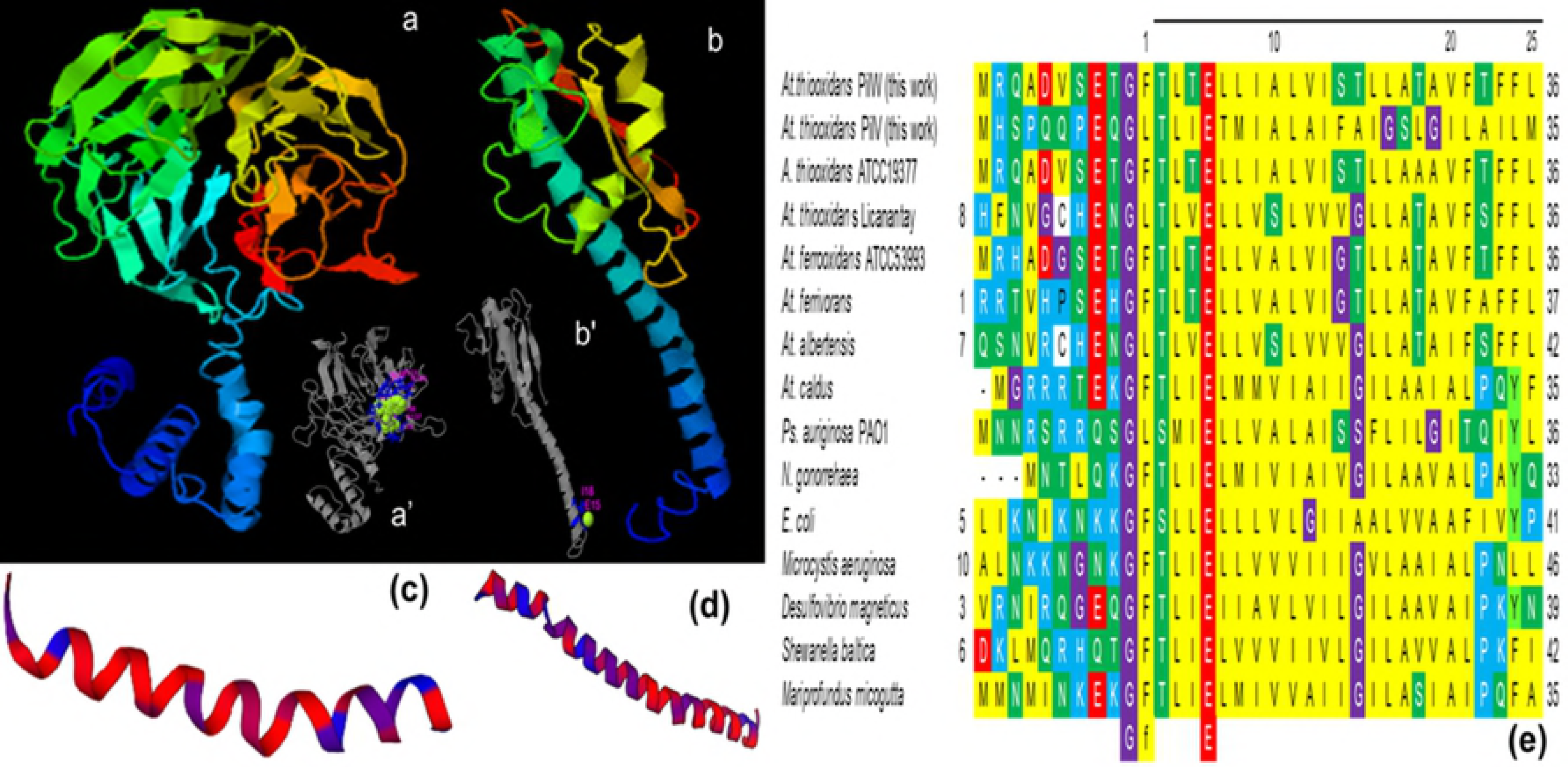
Phylogenetic relationships based on the pilins PilW (AWP39906.1) and PilV (AWP39907.1) of *At. thiooxidans*. The evolutionary history obtained by ML (log likelihood −1678.78) using 12 amino acids sequences. There was a total of 97 positions in the final dataset. Evolutionary analyses were conducted in [42]. The aa sequences used were derived from: *At. caldus* ATCC 51756 (WP_014002880.1); *At. ferrivorans* SS3 for PilW (WP_035191506.1), *At. ferrivorans* CF27 (CDQ09112.1), *At. ferrooxidans* (ACH83326.1), *At. thiooxidans* ATCC 19377 for PilW (WP_010638979.1) and PilV (WP_010638981.1), *Ps. aeruginosa* PilW, (NP_253242.1) and PilV (NP_253241.1) and *T. thermophilus* (AAM55484.1).

Resuming, PilY1 (AWP39905.1) comprises the conserved *Neisseria*-PilC (superfamily) beta-propeller domain of the tip-associated adhesin while PilW (AWP39906.1) of the superfamily of putative TfP assembly proteins, and PilV (AWP39907.1) belongs to the super family of TfP assembly protein. Further analysis will be required to elucidate the specific function of PilY1, PilW and PilV as well as the molecular mechanisms of pili assembly in *At. thiooxidans*.

## Acknowledgments

We thank LINAN-IPICyT, for providing laboratory support.

## Supporting information

**S1 Table**

**S2 Table**

## Authors’ contributions

JVGM and JAMC conceived and designed the experiments. EAS and AHS performed, described and discuss the sequencing. OASP conceived, designed, performed, described and discuss the TEM analyses. JVGM performed and discussed the bioinformatics analysis. MAG discussed the overall results. JAMC, JVGM and OASP contributed with reagents/materials/analysis tools. JVGM is senior author.

